# Impact of carbamate and pyrethroid pesticides on bile acid profiles in an in vitro gut microbiota model

**DOI:** 10.1101/2024.09.26.615257

**Authors:** Weijia Zheng, Wouter Bakker, Maojun Jin, Jing Wang, Ivonne M.C.M. Rietjens

## Abstract

The impact on the microbial community following exposure to commonly used pesticides has recently gained increasing interest. In this study, effects of selected carbamates (carbofuran, aldicarb) and pyrethroids (cypermethrin and cyhalothrin) on gut microbiota and related bile acid metabolism were quantified using a 24 h in vitro fermentation model system with mice feces. The results obtained reveal a pesticide induced significant increase the ratio of secondary over primary bile acids, particularly resulting from the enrichment of β-muricholate (βMCA) accompanied by the depletion of ω-muricholate (ωMCA). Besides, the bacterial profile showed significantly increased richness of *Eggerthellaceae* after 24 h exposure to carbofuran and cyhalothrin, and the genera *Enterorhabdus* belonging to *Eggerthellaceae* was found to be highly correlated to the fecal bile acid profile. In conclusion, in an in vitro gut microbial model carbamates and pyrethroids caused alterations of the gut microbial community resulting in the modulation of bile acid transformation. This illustrates that the gut microbiota and its metabolism may be a novel target to consider in future pesticide safety evaluations.

## 1. Introduction

In the past decades, the demand for food has risen significantly in relation to the world population’s increase, and as a result pesticides are widely used worldwide to obtain better quality agricultural products and increase crop yields, bringing significant economic benefits but also increased health hazards (Jin et al., 2017). Carbamates and pyrethroids are two typical classes of insecticides which are frequently used to control pests in homes and agricultural crops (Rawn et al., 2006). The systemic N-methyl carbamate pesticides such as carbofuran and aldicarb (Fig. 1) are also extensively applied as nematicide and acaricide for agricultural, domestic and industrial purposes (Mishra et al., 2020).

**Fig. 1.**
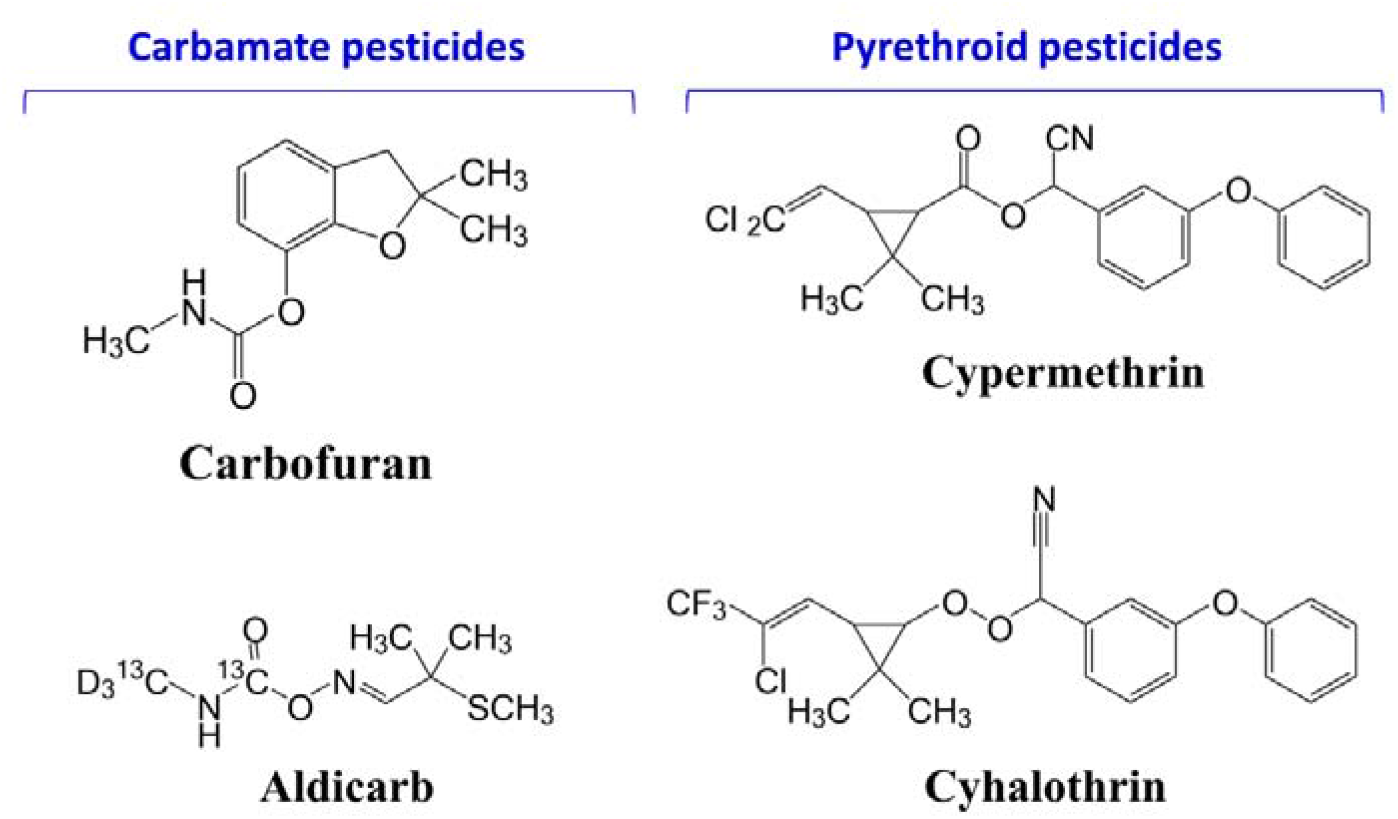
Chemical structures of the model pesticides of the present study including the carbamates carbofuran and aldicarb and the pyrethroids cypermethrin and cyhalothrin.

Furthermore, synthetic and effective pyrethroids like cypermethrin and cyhalothrin (Fig. 1) are largely replacing the use of organophosphorus insecticides due to the high risks of these organophosphorus to health and the environment (Muhamad et al., 2012). Recent work reported that residues of carbamates and pyrethroids in food may also cause neurotoxic, cytotoxic, reproductive, endocrine disrupting, embryo-toxic and dermal-skin problems (WHO, 2009). For example, it was indicated that oral administration of rats with carbamates could result in inhibition of cholinesterases (Silberman and Taylor, 2018), while accidental mammal exposure to pyrethroids could cause liver hypertrophy, mildly to severe irritation to skin and eyes, and neurotoxicity at high doses (Thatheyus and Selvam, 2013). Although regulation and control for use of carbamates and pyrethroids is being in place in many countries, it is inevitable that humans could be exposed to small amount of residues in a variety of food products (WHO, 2009), resulting particularly in the direct exposure of the gut and its microbiome. The gut microbiome consists of trillions of microorganisms that colonize the mammalian gastrointestinal tract. The past few years have seen a surge in research on the gut microbiome, which has firmly established its critical role in host health (Shreiner et al., 2015; Tremlett et al., 2017; Rooks and Garrett, 2016; Halfvarson et al., 2017). The gut microbiome is involved in numerous aspects of host metabolism and physiology, from energy production to stress response, and can confer many benefits to the host (Deehan and Walter, 2016; Dinan and Cryan, 2012). One important role relates to modulating lipid metabolism potentially through bacterial-derived signaling molecules such as bile acids (Ghazalpour et al., 2016). Bile acids are a class of structurally diverse molecules with a rich diversity and each of these entities may have different bioactive functions. In addition to their detergent-like properties and their use as substrates for microbial metabolism (Hylemon et al., 2018), bile acids act as hormones as well (Perino et al., 2021). As a result, the gut microbial community as well as the related bile acid profile play an important role in host health.

However, changes in the gut microbiota could be easily induced by various factors, for example, ageing (Leung and Thuret, 2015; Lynch et al., 2015), diet (Wu et al., 2016; Jeffery and O’Toole, 2013), diseases (Carding et al., 2015), and also exposure to chemicals including antibiotics, drugs and also some pesticides (Kang et al., 2013). Recently, the toxicity of pesticides towards non-target organisms such as the gut microbiota is gaining increased attention (Defois et al., 2018). Studies performed in vitro or in vivo on gut microbiota alterations and host outcomes induced by exposure to pesticides such as organochlorine pesticides, fungicides, and insecticides including carbamates have been reported (Giambò et al, 2021). In the only previous work evaluating the effects of carbamates on gut microbiota (Gao et al., 2018), the carbamate aldicarb showed effects on the gut microbiota including the increased abundance of *Erysipelotrichaceae* at the expense of *Christensenellaceae*, leading to changes in lipid profiling in C57BL/6J mice. The effects of pyrethroids including bifenthrin (Li et al., 2022; Li et al., 2021) and permethrin (Nasuti et al., 2016) causing gut microbiota dysbiosis have also been reported in recent years. However, scientific research for potential toxicity of cypermethrin, cyhalothrin, and carbofuran to gut bacteria and the related metabolome are still absent. To date, no studies have reported on the consequences of the effects of these pesticides on the microbiome for bile acid homeostasis.

The aim of the present study was to assess the potential effects of selected carbamate and pyrethroid pesticides on the gut microbiota and the related consequences for bile acid metabolism and the resulting bile acid profile. To this end, an in vitro model was employed enabling studying of the influence of selected carbamates and pyrethroids on mouse intestinal microbiota and the resulting effects on processing of bile acids. The in vitro model consisted of anaerobic fecal incubations, in which altered bile acid profiles could be quantified using liquid chromatography tandem mass spectrometry (LC-MS/MS) and bacterial profiles were determined by 16S rRNA gene sequencing analysis. Characterization of the effects of the exposure to carbamates and pyrethroids on the composition of gut microbiota and related bile acid metabolism will reveal whether the understanding of the interplay between oral administration of pesticides, gut microbiota and bile acid metabolism is an area of particular interest for future safety evaluations.

## 2. Materials and methods

### 2.1. Reagents and standards

Bile acid standards of taurocholic acid (TCA), tauro-β-muricholate (TβMCA), tauro-α-muricholate (TαMCA), taurodeoxycholic acid (TDCA), taurolithocholic acid (TLCA), tauroursodeoxycholic acid (TUDCA), taurohyodeoxycholic acid (THDCA), tauro-ω-muricholate (TωMCA), cholic acid (CA), β-muricholate (βMCA), α-muricholate (αMCA), ω-muricholate (ωMCA), deoxycholic acid (DCA), lithocholic acid (LCA), ursodeoxycholic acid (UDCA), and hyodeoxycholic acid (HDCA) were obtained from Merck KGaA (Darmstadt, Germany), or Cambridge Isotope Laboratories (Tewksbury, USA). The test pesticides carbofuran (CAS 1563-66-2), aldicarb (CAS 116-06-3), cypermethrin (CAS 52315-07-8), and cyhalothrin (CAS 68085-85-8), as well as dimethyl sulfoxide (DMSO) were purchased from Sigma Aldrich (Zwijndrecht, The Netherlands). Methanol and acetonitrile were supplied by Biosolve BV (Valkenswaard, The Netherlands), formic acid was ordered from VWR CHEMICA (Amsterdam, The Netherlands), and phosphate buffered saline (PBS) was purchased from Gibco (Paisley, UK). A mixture of taurine conjugated bile acids was prepared in PBS at final concentrations of 500 μM TCA and 500 μM TβMCA, and individual stock solutions of carbamates as well as pyrethroids were prepared in DMSO for further use.

### 2.2. Conversion of the in vivo dose levels to in vitro test concentrations

To establish the concentrations to be used in the in vitro studies, LD_50_ values reported for the respective pesticides in acute toxicity studies in mice were collected from the reports of the World Health Organization (WHO) and Joint Meeting on Pesticide Residues (JMPR). Since 1/10 LD50 is normally utilized as the low dose in toxicity studies of carbamate and pyrethroid pesticides, LD50 values of 30 mg/kg bw for carbofuran (WHO, 2003), 1.0 mg/kg bw for aldicarb (WHO, 2002), 250 mg/kg bw for cypermethrin (JMPR, 2006), and 20 mg/kg bw for cyhalothrin (JMPR, 2007) were converted to 3 mg/kg bw for carbofuran, 0.1 mg/kg bw for aldicarb, 25 mg/kg bw for cypermethrin and 2 mg/kg bw for cyhalothrin as the oral dose levels providing low but measurable effects without showing lethality. These dose levels were converted to the corresponding in vitro test concentrations as follows:

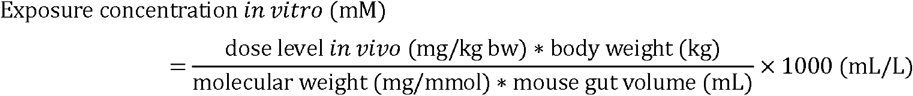

The volume of the gastrointestinal tract of mice was assumed to amount to approximately 1.5 mL (Davies and Morris, 1993) and the body weight of mouse was taken as 0.02 kg on average. Using the equation this resulted in test concentrations of 0.18 mM for carbofuran, 0.01 mM for aldicarb, 0.8 mM for cypermethrin, and 0.06 mM for cyhalothrin, concentrations that were employed as the test concentrations in the in vitro incubations.

### 2.3. Preparation of mouse fecal samples

Feces were obtained by physical massage of the rectum of C57BL/6N mice (30 male and 30 female mice), weighed and transferred immediately into anaerobic 10% (v/v) glycerol in PBS, pooled, homogenized and diluted to a final fecal concentration of 20% (w/v) under an anaerobic atmosphere (85% N_2_, 10% CO_2_, and 5% H_2_) in a BACTRON300 anaerobic chamber (Cornelius, USA). Samples were then filtered using sterile gauze under the anaerobic conditions, and 200 µL aliquots of fecal slurry were prepared. Subsequently, a washing step was implemented to avoid effects of high levels of the residual endogenous fecal bile acids in the developed model; as a result, these prepared aliquots were twice washed using an equal volume of anaerobic PBS, followed by vortex-mixing for 1 min and centrifugation at 2,000×g for 5 min at 4 °C under anaerobic conditions. Supernatants were removed and PBS was supplied to 200 µL ensuring that the washed fecal slurries contained 20% feces (v/v) as before. All the washed slurries were collected, mixed and newly aliquoted samples of 200 µL resulting fecal slurry were stored at -80°C until use.

### 2.4. In vitro incubations

Eppendorf tubes contained incubations consisting of 80 µL fecal slurry prepared in Section 2.3., 10 µL mixed solution of taurine conjugated bile acids (500 μM TCA and 500 μM TβMCA), and 10 µL DMSO (solvent control) or individual stock solution of pesticides. Thus, aliquots of 100 µL mixture with final concentrations of 50 μM TCA as well as 50 μM TβMCA, and 0.18 mM carbofuran or 0.01 mM aldicarb or 0.8 mM cypermethrin or 0.06 mM cyhalothrin with 16 % fecal slurry were incubated under anaerobic conditions. Incubations were performed in the BACTRON 300 anaerobic chamber (Sheldon, Cornelius, USA) with an atmosphere of 85% N_2,_ 10% CO_2_, and 5% H_2_, at 37 °C. At 0 h, 6 h, 12 h, and 24 h of this batch fermentation, the reaction was stopped by adding a similar volume (100 µL) of acetonitrile. Samples were subsequently sonicated for 5 min, centrifuged at 21500 g for 15 min at 4°C, and the supernatants obtained were stored at -80°C overnight, followed by freeze drying for 8h. Residuals obtained were then redissolved in methanol/water (1/1) acquiring the final volume of 100 µL for each aliquot. Subsequently, all samples were centrifuged at 21500 g for 15 min at 4°C, after which the supernatants were collected, filtered (PALL AcroPrep, PTFE 0.2 μm), and transferred to vials for measurement of bile acids by LC-MS/MS.

### 2.5. Bile acid profiling by LC-MS/MS analysis

Bile acid measurement was performed on a Nexera XR LC-20AD SR UPLC system coupled to a triple quadrupole LCMS 8050 mass spectrometer (Kyoto, Japan) with electrospray ionization (ESI) interface, which was able to measure the 16 bile acids studied herein: TCA, TβMCA, TαMCA, TCDCA, TDCA, TLCA, TUDCA, THDCA, TωMCA, CA, βMCA, αMCA, CDCA, ωMCA, DCA, LCA, UDCA, and HDCA.

Bile acids in fecal samples, fecal incubations and standards were separated on a Kinetex C18 column (1.7µm×100 A×50mm×2.1 mm; Phenomenex, Torrance, USA) using an ultra-high performance liquid chromatography (UHPLC) system (Shimadzu) with mobile phases consisting of 0.01% formic acid in distilled water (solvent A), a mixture of methanol and acetonitrile (v/v=1/1) (solvent B), and acetonitrile containing 0.1% formic acid (solvent C). The total run time was 16 minutes with the following gradient profile: 95% A, 0% B and 5% C (0-2min), slowly changed to 30% A, 70% B, and 0% C from 2 to 7.5 min, rapidly reversed to 2% A, 98 % B, and 0% C in 0.1 min, then kept at 2% A, 98% B, and 0% C from 7.6 to 10 min, then changed to 70% A, 30% B, and 0% C in 0.5 min; in the end, slowly turned back to the initial condition of 95% A, 0% B and 5% C from 10.5 to 13 min, then maintained at these conditions till 16 min for equilibration. The column temperature was set at 40°C and the sample tray temperature was set at 4°C. The mass spectrometer (MS) used electrospray ionization (ESI) in negative ion mode. The ESI parameters were as below: Nebulizing gas flow, 3L/min; drying gas flow and heating gas flow, 10L/min; Interface temperature, 300°C; Desolvation temperature, 526°C; heat block temperature, 400°C. Selective ion monitoring (SIM) as well as multiple reaction monitoring (MRM) were simultaneously used for the detection of the bile acids. Precursor and product ions were as flows: 407.3>407.3 *m/z* for αMCA, βMCA, ωMCA, and CA; 319.3>319.3 *m/z* for UDCA, HDCA, CDCA, and DCA; 498.4>498.4 *m/z* for TUDCA, THDCA, TCDCA, and TDCA; 514.4>514.4 *m/z* for TαMCA, TβMCA, TωMCA, and TCA; 375.3>375.3 *m/z* for LCA; 482.3>482.3 *m/z* for TLCA. The Postrun and Browser Analysis function from the LabSolutions software (Shimadzu, Kyoto, Japan) was employed to obtain the peak areas of the satisfied extracted ion chromatogram (EIC) for each target.

### 2.6. 16S rRNA gene sequencing analysis

Based on the results of bile acid profiling (see Section 3.2), carbofuran and cyhalothrin were respectively selected as the representatives of carbamate and pyrethroid pesticides for further study of the effects on the microbial community by 16s rRNA analysis. To this end, mixtures comprising 240 µL prepared mouse fecal slurry (Section 2.3) (final concentration 16 % feces) and 30 µL PBS, mixed with 30 µL DMSO (control) or 30 µL solution of pesticides (final concentrations 0.18 mM carbofuran or 0.06 mM cyhalothrin) were incubated under anerobic conditions for 24 h at 37°C. After 24 h, the fecal samples with or without pesticides were stored at -80°C overnight and subsequently delivered to an accredited commercial laboratory (IMGM Laboratories GmbH, Martinsried, Germany) for DNA extraction, PCR, library preparation, and sequencing. Besides, quantification of the bacterial load was implemented by real-time qPCR. 16S V3-V4 primers (F-NXT-Bakt-341F: 5′-CCTACGGGNGGCWGCAG-3′ and R-NXT-Bakt-805R: 5′-GACTACHVGGGTATCTAATCC-3′) were used to amplify the PCR products. During an index PCR, barcodes for multiplexed sequencing were introduced using overhang tags. A sequencing library was prepared from barcoded PCR products and sequenced on the Illumuna^R^ MiSeq next generation sequencing system (Illumuna^R^ Inc.). Signals were processed to *.fastq-files and the resulting 2×250 bp reads were demultiplexed. Microbiota identification was performed by clustering the operational taxonomic units (OTUs).

### 2.7. Data analysis

Metabolic profile data acquisition and processing were implemented using the Labsolutions software in the LC-MS/MS system. Graphics were drawn using Graphpad Prism 5 (San Diego, USA). Chemical structures were drawn by using ChemDraw 18.0 (PerkinElmer, Waltham, USA). Results are shown as mean ± standard deviation (SD) of three independent measurements and statistical significance was determined by a one-way ANOVA with a Dunnett/ Bonferroni correction for multiple tests and results were considered significant when p<0.05. 16S rRNA analysis data were analyzed with R version 3.6.1 and QIIME 2 view.

## 3. Results

### 3.1. Initial bile acid profile in the in vitro model

Fig. 2 presents the initial bile acid profile at 0 h of incubation, including levels of all quantified bile acids consisting of conjugated bile acids, primary bile acids, and secondary bile acids. Fig. 2A shows that following the optimized process of fresh isolation, filtration, washing with PBS, and addition of a mixture of taurine conjugated bile acids (final concentration of 50 µM TCA and 50 µM TβMCA), at t=0 h approximately 40 µM of TCA and TβMCA were detected in the samples together with a small amount of TαMCA detected as well. TCA and TβMCA were added to the incubations to better mimic intestinal bile acid profiles given that these conjugated bile acids are known to be excreted into the intestine from the liver, where they are present at high concentrations (Sayin et al., 2013). The primary and secondary BAs detected at t=0 mainly originate from the fecal samples, not being fully eliminated upon the washing procedure. This procedure of washing the fecal samples and adding TCA and TβMCA results in an initial bile acid profile that mimics the bile acid profile as excreted from the liver into the intestine (Sayin et al., 2013). The total amount of primary and secondary bile acids was quantified at a level of approximately 40 µM each. In Fig. 2B, it is shown that at the start of the incubations the conjugated and primary bile acids together make up of 83% of the overall bile acid profile, with the conjugated bile acids amounting to 50%. Secondary bile acids including ωMCA, DCA, LCA, and HDCA were quantified as well, and were present at concentrations below 10 µM each, together making up less than 20% of the total bile acid profile at the start of the incubations.

**Fig. 2.**
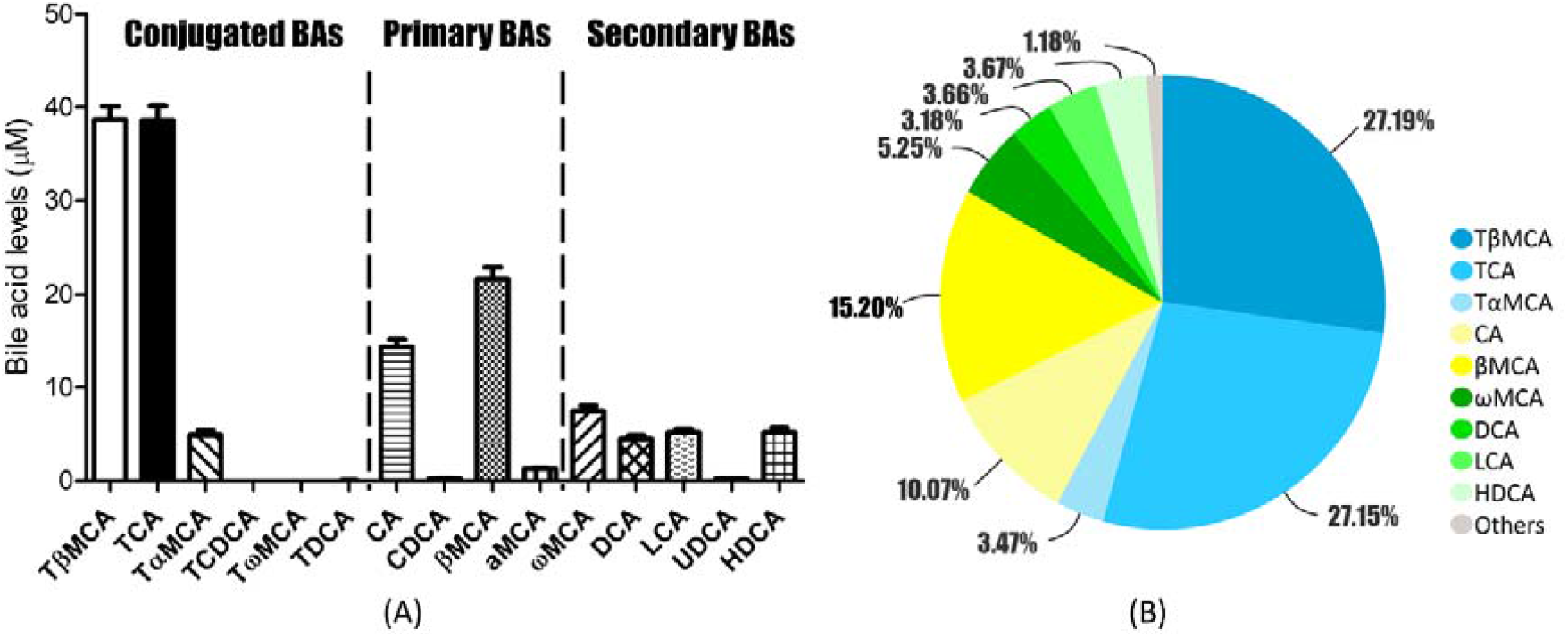
Initial bile acid profile at 0 h of incubation (n=3) represented as (A) absolute concentrations of the bile acids quantified including the conjugated bile acids TβMCA, TCA, TαMCA, TCDCA, TωMCA, and TDCA, the primary bile acids CA, CDCA, βMCA, and αMCA as well as the secondary bile acids ωMCA, DCA, LCA, UDCA, and HDCA and as (B) the relative percentage of the total bile acid profile showing the conjugated bile acids (blue), primary bile acids (yellow), secondary bile acids (green), and other bile acids at low concentrations (grey).

### 3.2. Bile acid profiles following 24 h incubation

Fig. 3 presents bile acid profile characteristics detected during the 24 h of the in vitro fecal incubations in the absence (solvent control) or presence of the selected pesticides. As an example, Fig. 3A shows the LC-MS chromatogram (obtained at precursor and product ions: 407.3>407.3) of the control and cyhalothrin treated incubation at 24 h, clearly revealing an increase in the amount of ωMCA and a reduced peak height of βMCA together with a stable peak height of CA. Fig. 3B presents the changes in the concentrations of the added taurine bile acids TCA and TβMCA during the 24 h anaerobic incubations; as observed, already upon 6 hours of incubation with or without pesticides, TCA as well as TβMCA were almost fully converted, reflecting fast deconjugation by the microbiota. Changes for other main fecal bile acids detected during the 24 h incubations are presented in Fig.S1, and the fecal bile acid profiles resulting at 24 h of incubation in the pesticide-treated samples and controls are shown in Fig. 3C. Significantly increased ωMCA and decreased βMCA levels were observed at 24 h in all pesticide-treated samples compared to controls, fully in line with the chromatogram shown in Fig. 3A. In addition, significantly decreased concentrations of αMCA were observed in carbofuran-, cypermethrin- and cyhalothrin-treated fecal incubations, while significantly reduced levels of DCA were observed in only the carbofuran and cyhalothrin groups compared to controls. Besides, at 24 h UDCA was detected at low concentrations <0.25 µM in all the pesticide-treated and control samples with a small but significant increase detected only in cypermethrin-treated samples (p<0.05). No significant changes were found at 24 h of incubation in the levels of CA, LCA, and HDCA when comparing results from the pesticide-treated samples and controls. In addition to the evaluation of changes in the individual bile acids, the data were also analysed with respect to two important characteristics of bile acid profiles, including the proportion of conjugated bile acids (%conjugated BAs) and the ratio of secondary bile acids to primary bile acids (secondary/ primary BAs). Fig. 3D presents the results obtained and reveals that the ratio of secondary/ primary BAs was significantly increased in all the pesticide-treated samples compared to controls, whereas significant changes in the % conjugated BAs were only observed in the pyrethroid-treated samples, showing decreases upon incubation with cypermethrin and cyhalothrin as compared to the control. Moreover in Fig. 3E, the results of a principal coordinates analysis (PCoA) of the bile acid profiles obtained from all fecal samples is presented. It shows that the controls as well as all pesticide-treated samples cluster in their own groups, with all the pesticide groups clearly separating from the controls.

**Fig. 3.**
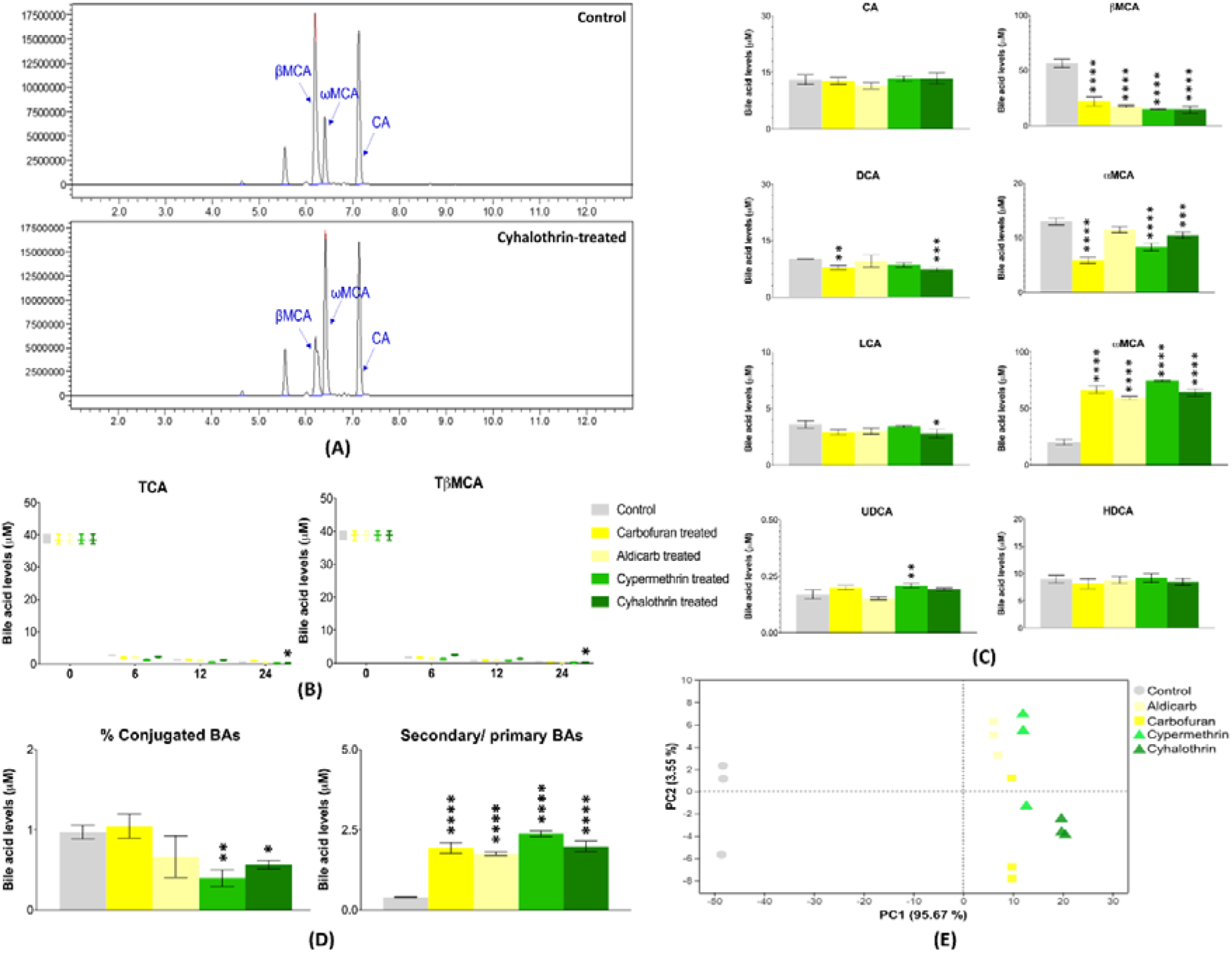
The altered bile acid profiles in the carbofuran, aldicarb, cypermethrin, or cyhalothrin -treated samples and controls, including (A) the chromatography of 407.3>407.3 m/z showing the altered peaks of βMCA, ωMCA, but not for CA in control and cyhalothrin (as example of pesticides) -treated fecal samples at 24 h of incubation, (B) the degradation of the added TCA and TβMCA at 6, 12, and 24 h following the incubations, (C) the changes of individual fecal bile acids detected in the pesticide-treated and control samples at 24 h of incubations, and (D) the altered bile acid profiles when expressed as the proportion of fecal conjugated bile acids (% conjugated BAs) and the ratio for the total amount of fecal secondary bile acids to primary bile acids (secondary/ primary BAs) (*p<0.05, **p<0.01, ***p<0.001, ****p<0.0001; n=3). Moreover, (E) shows a principal coordinates analysis (PCoA) performed based on the bile acid profiles in the pesticide-treated and untreated fecal samples.

To facilitate interpretation of the results the metabolic pathways for the detected individual bile acids including conjugated, primary, and secondary bile acids is presented in Fig. 4. The main bile acids detected can be divided into two groups, composed of the bile acids originating from the added TCA or from the added TβMCA. Combining these pathways with the results shown in Fig.3, reveals that the bile acids in the TβMCA pathway showed more significant alterations than those from the TCA based pathway. The combined data also indicate that conversions like bile salt hydrolase (BSH) deconjugation, epimerization, dehydroxylation, and hydroxylation were affected to a different extent upon the treatment of the fecal microbiota with different pesticides. The 6β-epimerization from βMCA to ωMCA was the most significantly affected conversion and this conversion was obviously accelerated upon the exposure to pesticides, leading to the raised levels of ωMCA at the cost of βMCA and an increase in the ratio secondary/ primary bile acids. Meanwhile, in pyrethroid-treated samples, the decreased % conjugated BAs indicated an increase in the BSH deconjugation.

**Fig. 4.**
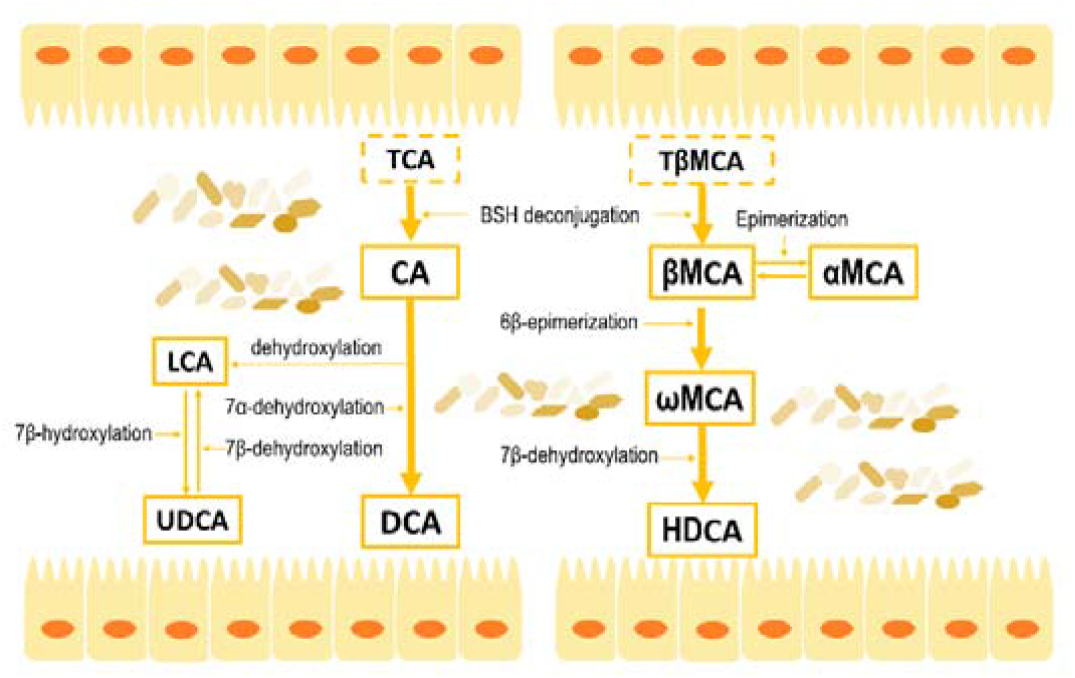
The bile acid transformation routes, starting from the added conjugated bile acids TCA and TβMCA, and including the formation of the primary bile acids CA, βMCA, and αMCA, and the secondary bile acids DCA, LCA, UDCA, ωMCA, and HDCA (Wahlström et al., 2016).

### 3.3. Alteration of the microbial community in anaerobic fecal incubations upon treatment with pesticides

Based on the results obtained for the bile acid profiles (Section 3.2), the carbamate carbofuran and the pyrethroid cyhalothrin were selected for further studies on their effects on the bacterial profiles, because these two pesticides resulted in the most pronounced effects on the bile acid profile. Fig. 5 presents the relative abundance of gut bacteria at phylum and family level as well as the analysis of alpha diversity in the fecal samples of the control and carbofuran or cyhalothrin treated groups. As shown in Fig. 5A, the phyla of *Bacteroidetes* and *Firmicutes* dominated in mice feces, followed by a small amount of *Actinobacteria*. The relative abundance of the phyla *Actinobacteria* significantly increased in carbofuran (p<0.01) and cyhalothrin (p<0.05) treated groups compared to controls, whereas the richness of the phyla *Bacteroidetes* was significantly reduced only in the carbofuran treated samples (p<0.05). At the family level (Fig. 5C), *Erysipelotrichaceae* belonging to *Firmicutes* and *Muribaculaceae* belonging to *Bacteroidetes* were the most abundant bacterial species, accounting for more than 50 % of the mouse gut microbial community. In addition, *Eggerthellaceae* was the most dominant family belonging to the phyla *Actinobacteria*, the relative abundance of which significantly increased in the carbofuran-treated (p<0.01) and cyhalothrin-treated (p<0.01) samples compared to controls. Besides, a raised richness of *Desulfovibrionaceae* was observed in both pesticide-treated samples especially in those exposed to cyhalothrin. The richness of *Anaeroplasmataceae* was reduced and the proportion of *Enterobacteriaceae* was increased in carbofuran-treated and cyhalothrin-treated groups, respectively, albeit not to a significant extent. Further, the alpha diversity based on the data OTUs is described in Fig. 5B utilizing the Chao1 and Shannon index, which separately indicate the total amount of bacterial species and the bacterial diversity. Results obtained show that the number of bacterial species as well as the diversity were reduced in fecal samples upon exposure to carbofuran, while both were slightly increased in the cyhalothrin-treated group, although these effects were not significant compared to the controls.

**Fig. 5.**
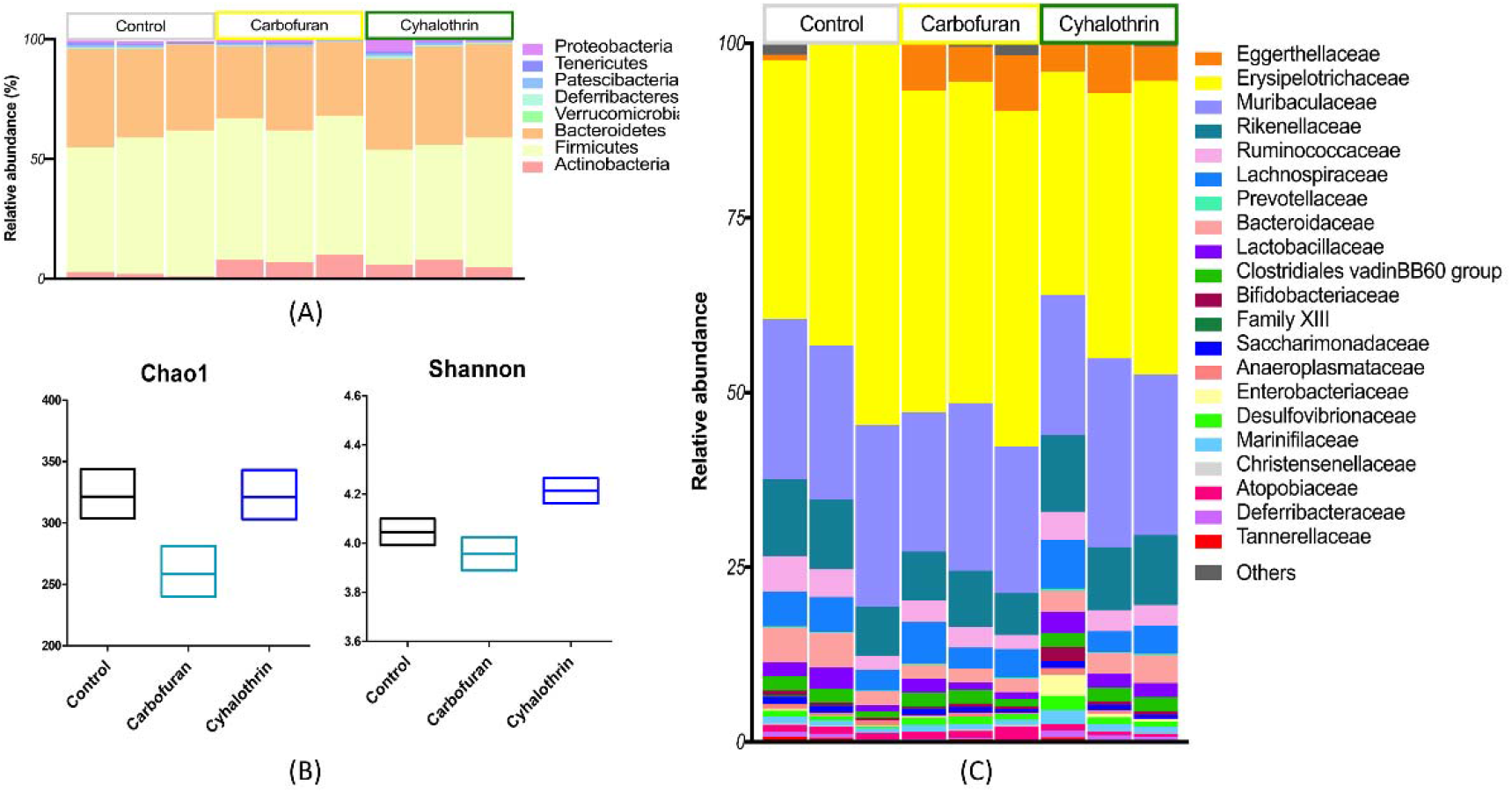
Composition of the microbiota community in untreated and pesticide treated fecal samples (n=3) after 24 h incubation in the absence (control) or presence of carbofuran or cyhalothrin, at (A) phylum level and (C) family level, and (B) the alpha diversity calculated based on the data of operational taxonomic units (OTUs) and shown by the index of Chao1 and Shannon.

### 3.4. Correlation of bile acid profile and microbiota

Fig. 6A presents the results of a pearson correlation analysis to identify the relation between changes in the microbial community and the altered bile acid profiles upon 24 h incubation in carbofuran-treated and cyhalothrin-treated fecal samples, respectively. In general, the family of *Eggerthellaceae* showed to be the most highly related to the fecal bile acids, being positively correlated with ωMCA and negatively correlated with the primary bile acids βMCA and αMCA; additionally, it was also negatively related to TCA in the carbofuran-treated samples. Another family *Desulfovibrionaceae* also presented strong correlations with fecal bile acid profiles, showing a negative association with TCA in the carbofuran group, and with TβMCA as well as LCA in the cyhalothrin group, and with αMCA in both pesticide-treated samples. Besides, *Anaeroplasmataceae* presented a highly negative correlation with CA in the carbofuran group, whereas *Enterobacteriaceae* appeared to be negatively correlated to UDCA in the cyhalothrin group. All the detected bacterial species involved in the four families that showed a link at genus level with the bile acid profiles are summarized in Fig. 6B. As shown, the genera *Enterorhabdus* (>5 % of the bacterial community), *Desulfovibrio* (>1 % of the bacterial community), *Anaeroplasma* (<1 % of the bacterial community), *Escherichia-Shigella* (<1 % of the bacterial community) were the dominant bacterial species in the families *Eggerthellaceae, Desulfovibrionaceae, Anaeroplasmataceae*, and *Enterobacteriaceae*. In the pesticide-treated samples, the richness of *Enterorhabdus* significantly increased (0.001<p<0.01 in carbofuran; 0.01<p<0.05 in cyhalothrin) at the expense of *Desulfovibrio*; additionally, the relative abundance of *Anaeroplasma* decreased in carbofuran-treated samples and that of *Escherichia-Shigella* markedly increased in cyhalothrin-treated samples. *Enterorhabdus* as well as *Desulfovibrio* appeared to be highly correlated to the bile acid profiles in both pesticide-treated samples. Further in Fig. 6C, a linear regression model was utilized to reflect the correlation of these genera with the two crucial bile acid profile parameters including the % conjugated BAs and the ratio secondary/ primary BAs (Fig. 3D) and the R square values were calculated. As shown, the genera *Enterorhabdus* was found to have a significant and strong correlation with the fecal ratio secondary/ primary BAs (R^2^=0.7).

**Fig. 6.**
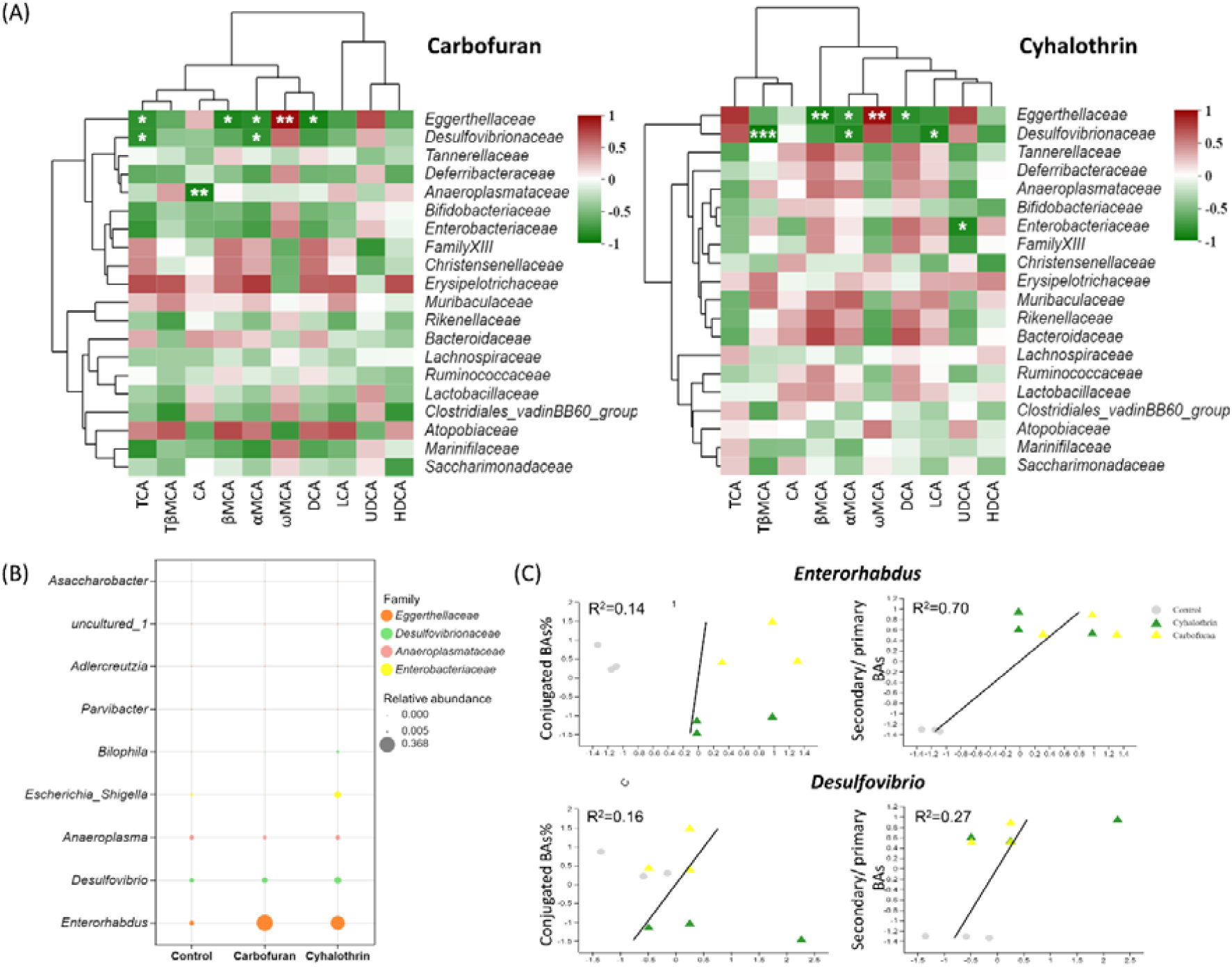
Correlations between the alterations within the bacterial community (family level) and the changes in bile acid profiles in carbofuran- and cyhalothrin-treated samples as reflected by (A) hierarchic clustering indicating statistical significance of the correlations found, (B) a bubble diagram presenting the changed richness of genera at the family level that showed high correlation with bile acid profiles and (C) linear regression plots of the correlations of the two mostly correlated genera shown in (B) with the bile acid profiles.

## 4. Discussion

The gut microbiota, in addition to their ability to process dietary derived material, are also capable of performing a range of bio-transformations on xenobiotics such as drugs and their metabolites; on the other hand, the gut microbiota can be affected by these compounds as well. In recent years studies on effects on the gut microbiota by xenobiotics, including for example antibiotics (Lange et al., 2016), antidiabetic drugs (Montandon and Jornayvaz, 2017) or some environmental pollutants including pesticides (Jin et al., 2017) have been reported. Particularly, some of the widely used pesticides were found to be able to alter the microbiota composition, which was suggested to lead to colon diseases and liver failure (Jin et al., 2017). However, very few studies have been published on the consequences of pesticide exposure of the gut microbiota for the related metabolism and health. In the present study, using an in vitro anaerobic incubation model, we revealed, for the first time, that the exposure of the gut microbiota to selected carbamates and pyrethroids could modulate fecal bile acid profiles and additionally, it was shown that effects of carbofuran and cyhalothrin on the microbial community correlated with the alterations in bile acid profiles.

Results obtained for the altered bile acid profiles induced by the treatment of carbamates and pyrethroids (Fig. 3) together with the bile acid transformation routes shown in Fig. 4 revealed that the 6β-epimerization of βMCA to ωMCA was significantly increased. In Fig. 6, *Eggerthellaceae* simultaneously showed its negative correlation with βMCA and positive link to ωMCA; as a result, the raised richness of *Eggerthellaceae* could be correlated to the faster 6β-epimerization. Additionally, another epimerized metabolite of βMCA, αMCA was also reduced in its richness, which could originate from the decrease in βMCA and was also related to the increased abundance of *Eggerthellaceae*. In addition to the bile acid originating from the added TβMCA (Fig. 4), the ones originating from TCA were also altered albeit to a less significant extent than the ones originating from TβMCA. For example, reduced levels of DCA were observed in both carbofuran and cyhalothrin treated samples, which could also be associated with raised richness of *Eggerthellaceae. Eggerthellaceae* belongs to the phyla *Actinobacteria* which has been previously reported for its beneficial effects on the degradation of toxic chemicals such as pesticides (Shivlata and Satyanarayana, 2017); meanwhile, *Eggerthellaceae* as an important member of *Actinobacteria* has also been found for its specific function on steroid metabolism (Hylemon et al., 2018). At the genus level, the dominant member of *Eggerthellaceae, Enterorhabdus* showed a significantly high correlation with the fecal bile acid profiles (Fig. 6B and C) especially with the conversion of βMCA to ωMCA, thereby leading to an increased ratio of secondary/ primary bile acids. To date, few scientific research on *Enterorhabdus* has been published, however, it could be a biomarker for potential effects caused by exposure to xenobiotics. Some studies reported a significantly increased abundance of *Enterorhabdus* in case of rat small intestinal cell line IEC-18 cell damage (Chen et al., 2020), altered type of diet (Pagliai et al., 2020), and treatment with xylooligosaccharide (Li et al., 2015) in a mouse model. In the cyhalothrin group, the increased richness of *Desulfovibrionaceae* might have significantly raised the deconjugation of TβMCA leading to the obvious decrease of the % conjugated BAs at 24 h of incubation. Although *Desulfovibrionaceae* only makes up a small proportion in the mouse gut microbial community, it has been found to be related to the bile acid metabolism (Hu et al., 2022). Meanwhile, for the genera *Desulfovibrio* dominating in *Desulfovibrionaceae* its raised abundance has been associated by others with gallstone disease and colorectal cancer in human (Ajouz et al., 2014; Ou et al., 2012).

On the other hand, not only the gut microbiota act on the bile acids, the altered bile acid profiles could also modulate the microbial community. For example, the conjugated bile acids were previously found to exhibit antibacterial activity (Jones et al., 2008; Inagaki et al., 2006); as a result, the observed reduced bacterial diversity in the carbofuran group at 24 h (Fig. 5B) was shown to be related to the slightly increased proportion of conjugated bile acids (Fig. 3D). Additionally, it has been reported that the ratio of secondary/ primary bile acids can be positively correlated to certain commensal genera (Kakiyama et al., 2013), hence the raised richness of *Enterorhabdus* may be ascribed to the increased ratio of secondary/ primary bile acids induced by the exposure to pesticides as well.

Collectively it can be concluded that, using the in vitro fermentation model it was shown that the exposure to pesticides significantly altered the gut microbial community resulting in changes of related bile acid profiles. A large amount of work has reported on the crucial role of bile acid signaling for host health (Perino et al., 2021); however to our knowledge, there is only one published study which revealed changes of gut microbiota-related bile acid profiles upon the exposure to pesticides. This study reported that the exposure to organochlorine pesticides induced an enhanced abundance of the genera *Lactobacillus* with BSH activity leading to altered bile acid metabolism. Thus, the present study provided a new proof for the effects of pesticides on gut microbiota and related bile acid metabolism using an in vitro model. To what extent the in vitro elucidated changes in the microbiome pattern and bile acid homeostasis are to be expected in the in vivo situation remains of interest for future studies.

## 5. Conclusion

In the current study, effects on gut microbiota and related bile acid metabolism caused by the exposure of mice fecal microbiota to carbamate and pyrethroids pesticides including carbofuran, aldicarb, cypermethrin and cyhalothrin were evaluated using an optimized in vitro fermentation model system with mice feces. Results obtained revealed a significant enrichment of secondary bile acids especially ωMCA with an accompanying decrease of the primary bile acid βMCA, indicating that the exposure of the microbiota to carbamate and pyrethroids could induce a faster transformation from primary bile acids to secondary bile acids. Meanwhile, an increased abundance of the bacterial family *Eggerthellaceae*, dominant in the phyla *Actinobacteria*, was observed in carbofuran and cyhalothrin treated samples; additionally, *Enterorhabdus* belonging to the *Eggerthellaceae* was found to be highly correlated to the fecal bile acid profiles. Our findings contribute to the issue on studying the altered gut microbial community and its related metabolism induced by pesticides, and point at potential effects on the host–microbial relationship upon pesticide exposure. The results also illustrate that the gut microbiota and its metabolism could be a novel target to consider in future pesticide safety evaluations.

## Acknowledgements

The authors gratefully acknowledge prof. dr. Jacques Vervoort for his supervision and efforts during the work. We also acknowledge Bert Spenkelink for help with collection of mouse fecal samples.

We thank the fundings provided by Key Lab of Agro-product Quality and Safety (Beijing, China).

---

**Fig. S1.**
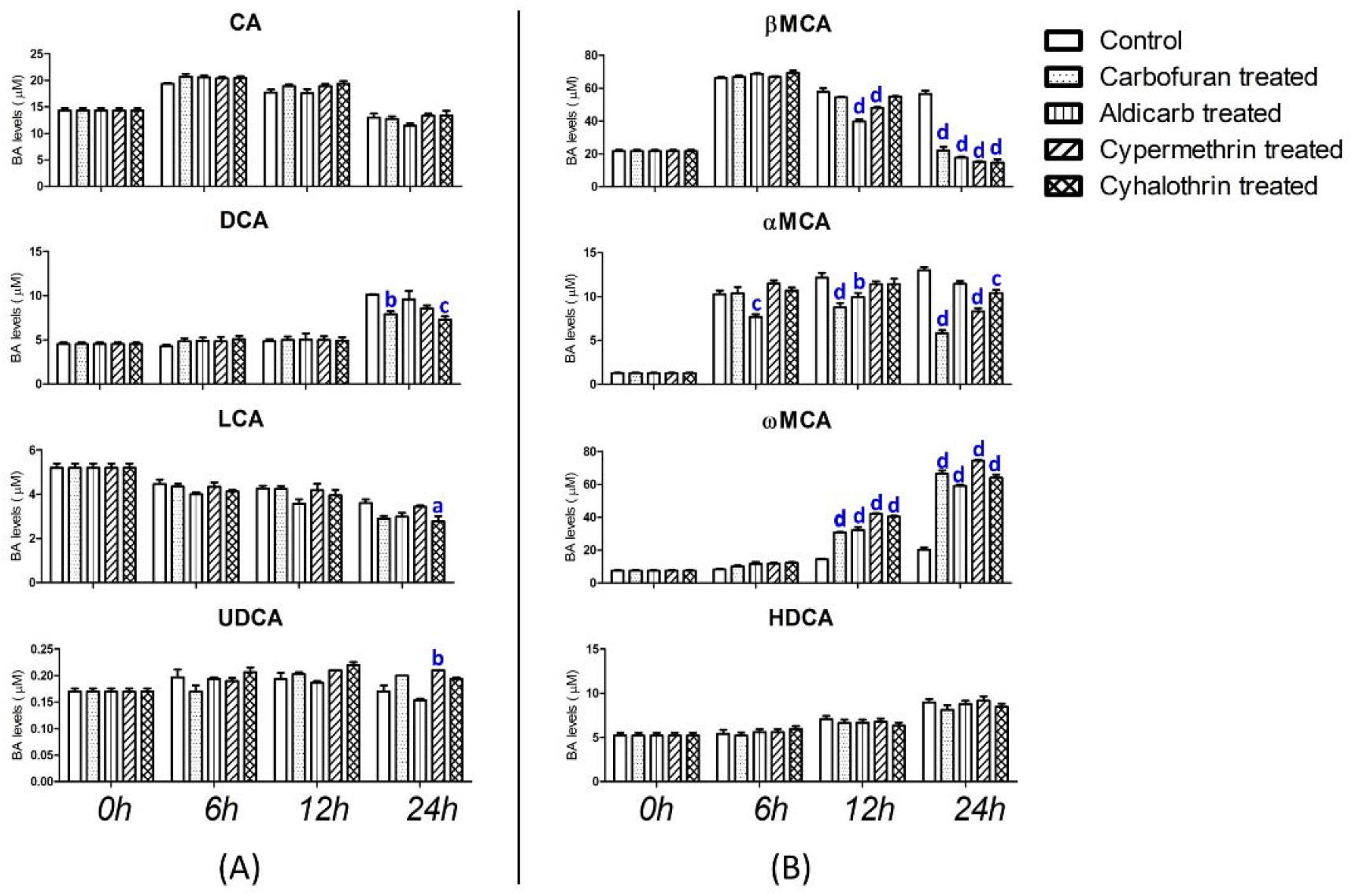
Bile acid changes in in vitro anaerobic mouse fecal incubations with or without (control) treatment with carbamates (carbofuran and aldicarb) and pyrethroids (cypermethrin and cyhalothrin) following the 24 h incubations (n=3). Transformation of (A) the added TCA, and the resulting changes in TCA, CA, DCA, LCA, and UDCA, as well as (B) the added TβMCA and the resulting changes in βMCA, αMCA, ωMCA, and HDCA. Samples were taken at 0 h, 6 h, 12 h, and 24 h during the incubation, and significance was marked using a (*p<0.05), b (**p<0.01), c (***p<0.001), d (****p<0.0001), which indicates a difference with control.

